# Engineering macrophage responses through 3D scaffold microarchitecture

**DOI:** 10.1101/2025.07.16.665133

**Authors:** Chiara Martinelli, Srijan Chakraborty, Giovanni Buccioli, Matteo Vicini, Claudio Conci, Giulio Cerullo, Roberto Osellame, Giuseppe Chirico, Emanuela Jacchetti, Manuela T. Raimondi

## Abstract

Biomaterial implantation in living organisms triggers a physiological response known as foreign body reaction, leading to the recruitment of macrophages, that can polarize either into a pro-inflammatory (M1) or an anti-inflammatory (M2) phenotype. Currently, there is growing interest in tailoring the physical properties of tissues and biomaterials to promote efficient tissue regeneration. Tridimensionality can profoundly influence macrophage behaviour; however, there is no clear consensus on the underlying mechanisms. 3D microstructures may play a crucial role in modulating immune cells, promoting anti-inflammatory responses, and supporting effective tissue repair and regeneration. In this study, we used two-photon polymerization to fabricate 3D scaffolds with large pores, measuring 50x50x20 μm³, and small pores, measuring 15x15x15 μm^3^. Both microstructures effectively influenced macrophage cytoskeletal organization and cellular metabolic activity. Notably, they were not sufficient to induce spontaneous macrophage polarization, indicating that they are intrinsically immunologically inert. When combined with chemical stimulation, as typically occurs *in vivo*, they elicited distinct responses. Specifically, as evidenced by the slight upregulation of the Arg1 marker, large pore sizes promoted an anti-inflammatory phenotype. Conversely, iNOS expression measurements indicated that small pores, which impose spatial constraints on macrophages, favoured a massive pro-inflammatory state. Our results demonstrate that 3D microstructures are versatile tools for multiple applications. Their precisely tunable architecture enables fine control over macrophage behaviour and immunomodulation, opening new avenues both for tissue engineering, by preventing fibrosis and promoting anti-inflammatory and pro-regenerative responses *in vivo*, and for the development of *in vitro* platforms to model inflamed tissues for screening anti-inflammatory drugs.

## 1. Introduction

Currently, the global market for implantable medical devices is valued at US $ 131.9 billion in 2024 and is expected to reach a value of nearly US $ 306.8 billion by 2034 [1]. Devices based on biomaterial implantation in living organisms trigger a physiological response, called foreign body response (FBR), characterized by the involvement of several cellular populations and biological components triggering a cascade of events crucial in determining their fate [2,3]. Generally, excessive chronic inflammation and fibrosis affect the device’s functionality and durability. Considering a constant failure rate of 10 % for all implantable devices due to FBR, the value of addressing this critical clinical need is worth US $10 billion per year. Moreover, medical device-associated infections represent a relevant concern both at the clinical and economic levels. Indeed, they remain difficult to be treated, with high risk of complications and many of them require device removal and antibiotic administration over long time periods. Approximately 60-70 % of healthcare associated infections are linked to medical device usage, and, in the US, it has been estimated that their rate is around 6.6 % [4]. According to the report by World Health Organization (WHO), healthcare associated infections range between 3.6 and 12 % in high-income countries and up to 19.1 % in low- and middle-income countries [5]. Understanding how to modulate FBR and avoid device-related infections is essential for achieving a successful reparative response [6]. During the entire wound healing process, immune cells play a fundamental role in modulating the progression of inflammation [3]. Macrophages are highly plastic cells, specialized in phagocytosis and able to secrete cytokines stimulating the proliferation and differentiation of the monocytic population [2]. They can polarize into i) a pro-inflammatory phenotype (M1), which favours inflammation by releasing pro-inflammatory cytokines and presents a phagocytic activity, necessary to remove bacteria and debris from the site of implantation, or ii) an anti-inflammatory phenotype (M2), which promotes tissue healing and angiogenesis through the secretion of anti-inflammatory cytokines and growth factors [7]. These cells are fundamentally involved in balancing between inflammatory and regenerative outcomes, through the release of definite molecules and growth factors, and determine whether the device will undergo integration, with tissue repair and regeneration, or rejection, with massive fibrous encapsulation [8]. M1 macrophages are crucial to the early pro-inflammatory response; however, their prolonged presence can lead to severe FBR. Conversely, excessive persistence of M2 macrophages can lead to an unresolved regeneration phase with thick fibrous capsule formation, and often results in unsuccessful biomaterial integration [9]. The balance between the two phenotypes is reversible and the co-existence of both phenotypes in the same microenvironment can be guided to modulate the inflammatory reaction [9]. Since macrophages have the tendency to quickly adapt to the surrounding stimuli, their activation is a complex process, influenced by the temporal and spatial presence of cytokines, other cells and physical signals [10]. Indeed, macrophages are susceptible to mechanical cues and manipulating the physical and mechanical properties of biomaterials can be considered the most promising strategy for achieving an optimal control of their phenotype, directing them toward an anti-inflammatory response [11]. Nowadays, there is growing interest in tailoring the physical properties of tissues and biomaterials to enhance efficient tissue regeneration, thanks to the possibility of limiting pro-inflammatory responses. Research in this field has been focusing on the 2D features of biomaterials, such as surface topography (*i.e.,* roughness, alignment and pattern of the surfaces) [13–16] and on 3D architecture (*i.e.,* shape, porosity) [10, 12, 17]. While 2D materials allow unravelling behaviour of macrophages in response to an external stimulus in simplified models, their physiological responses can be better appreciated in more realistic 3D microenvironments. Fabricated 3D microstructures play a crucial role in modulating immune cells and a rationale design of the geometry of implantable devices can be envisaged for promoting anti-inflammatory responses, thereby supporting more effective *in situ* regeneration. There is a strict correlation between the physical properties of the 3D microenvironment and the efficiency of tissue repair and regeneration. In general, an increase in pore diameter enhances nutrient transport and oxygen diffusion, facilitates waste removal, and promotes cell infiltration, proliferation, and vascularization [6]. However, a clear consensus on the optimal pore size for directing macrophage behaviour towards inflammation resolution, tissue remodelling, and effective biomaterial integration is still lacking. This gap in knowledge is partially due to the limitations of current biomaterial fabrication technologies, which focus primarily on modulating the mechanical properties rather than precisely controlling pore architecture. Currently, within the global scientific landscape, there is a gap in the investigation of this phenomenon at certain dimensional scales. In fact, many studies focus on the surface topography of grafts and implants at the nano- and sub-micrometer scales [13–16], while another research branch utilizes mesoscale scaffolds [18–20], with pore sizes in the order of hundreds of micrometers, primarily aiming to optimize tissue-level integration. Interestingly, only a few investigations have addressed intermediate pore sizes, in the range of tens of micrometers, which are particularly relevant because they approximate the size of individual cells or small cell clusters. For example, Madden *et al.* observed, upon cardiac implantation of poly(2-hydroxyethyl methacrylate-co-methacrylic acid) hydrogel scaffolds with pore size of 30-40 μm, increased angiogenesis and reduced fibrosis, favoured by a transition of macrophages towards the M2 phenotype [21]. Alternatively, macroporous polycaprolactone scaffolds, with fiber diameters of 5-6 μm and pore sizes of approximately 30 μm, have been investigated as potential vascular grafts. These scaffolds promoted cell infiltration and extracellular matrix deposition, and induced the recruitment of M2-polarized macrophages, thereby supporting artery regeneration and vascularization [22]. In addition, using higher-resolution scaffolds composed of electrospun polydioxanone fibers with diameters progressively increasing up to 2.8 ± 0.5 μm, and pore radius reaching 14.73 ± 0.72 μm, Garg *et al.* observed that bone marrow derived macrophages polarized *in vitro* preferentially expressed the Arginase 1 (Arg1) biomarker, specific to the M2 phenotype, showing a clear correlation with scaffold architecture [23].

In the present work, we report a radically novel approach for mimicking the physiological tissue microenvironment *in vitro*, by analysing macrophage polarization phenotypes in presence of both microstructured substrates and chemical induction. Our engineered microstructures align with the above-mentioned research, focused on investigating the microenvironment at the cellular scale, crucial for understanding and modulating immune cell behaviour in tissue regeneration contexts. Since it is well established that the earliest cellular responses to the surrounding microenvironment involve both cytoskeletal remodelling and metabolic reprogramming, which ultimately guide the cell toward its specific differentiation fate [11, 24–26], in this study we conducted an in-depth analysis of these two key aspects. Specifically, we quantitatively evaluated a set of morphological parameters related to filamentous actin (F-actin) organization, which reflect cytoskeletal architecture and mechanotransductive adaptation. In parallel, we assessed cellular metabolic activity using Fluorescence Lifetime Imaging Microscopy (FLIM), a powerful, label-free technique capable of detecting shifts in metabolic state based on the fluorescence lifetimes of endogenous cofactors such as nicotinamide adenine dinucleotide phosphate (NAD(P)H). Free (NAD(P)H) has shorter fluorescence lifetime, corresponding to increased glycolysis, while protein-bound (NAD(P)H) has significantly longer fluorescence lifetime, reflecting a shift of metabolism to oxidative phosphorylation [27–31]. In our work, FLIM has been exploited to classify the phenotypes of macrophages cultured in 2D and 3D microenvironments, with or without chemical treatments aimed at inducing M1 or M2 polarization, by analysing their metabolic profiles. Indeed, M1 macrophages primarily rely on glycolysis, while M2 macrophages mainly depend on oxidative phosphorylation [27, 28]. This metabolic state can be detected with the corresponding variations in the lifetimes of NAD(P)H inside the cell. The mean lifetime of the cells, calculated by the weighted sum of the lifetimes of free and protein bound NADH with their amount present inside the cells, is used to gauge their metabolism. A lower mean NAD(P)H fluorescence lifetime reflects a shift towards glycolytic metabolism, as seen in M1 macrophages [29–31].

## 2. Materials and methods

### 2.1 3D microstructure fabrication by two-photon laser polymerization

Two-photon fabrication was performed as previously published by our group [32–34]. Briefly, a Yb:KYW cavity-dumped laser source was used. This provided femtosecond pulses having λ = 1030 nm, repetition rate 1 MHz, pulse energy 1 μJ, pulse duration of ∼ 300 fs. The employed fabrication setup is based on three planar motion stages (ANT130XY Series and ANT130LZS, Aerotech, USA) controlled by a proprietary numerical control software, and on a spatial light modulator (SLM, PLUTO NIR-049, HOLOEYE, Germany). The laser beam was focused in the photoresist *via* a 100 x, NA = 1.4, oil immersion objective (Plan-Apochromat, Carl Zeiss, Oberkochen, Germany). The employed photoresist is the SZ2080 previously validated, in terms of optical quality, mechanical stability, biocompatibility and low shrinkage properties [35–40]. In this work, we dropcasted 46 μl of liquid resist on circular glass coverslips (⌀ = 12 mm, thickness = 170 μm, ThermoFisher Scientific, USA). Resist performed a pre-condensation at room temperature for 48 hours before its polymerization. A development procedure employed a solution of propan-2-ol and methyl isobutyl ketone (Sigma-Aldrich, USA) in a ratio of 50:50 (v/v) for removing the unpolymerized photoresist after the fabrication process (developing time: 35 minutes). Scanning Electron Microscopy (SEM, Phenom Pro, Phenom-World, Netherlands) was performed to check the quality of the fabricated samples. Two microstructures with different interconnected pores were designed and fabricated (**Figure 1**): one having 50x50x20 μm^3^ pores, referred to as 50x50, validated in a previous work [32], and the other having 15x15x15 μm^3^ pores, referred to as 15x15. The 50x50 scaffold is constituted by a total of 200 pores, with a height of 40 μm and a total dimension of 500x500x40 μm^3^. The 15x15 microstructures feature instead a lattice organization of two superimposed layers constituting a scaffold of a total of 784 pores, a height of 30 μm and a total dimension of 420x420x30 μm^3^. Both scaffolds were inserted into cell culture substrates (*i.e.,* 24-well plates) and sterilized for subsequent *in vitro* experiments. Imaging sessions were conducted in two configurations: in the first, the functionalized coverslip was glued to the bottom of plastic Petri dishes (Loctite 3943, Henkel) to perform FLIM acquisitions; in the second, after cell fixation, the functionalized glass coverslips were mounted onto rectangular microscope slides to perform confocal laser scanning microscopy. The microfabricated scaffolds for the *in vitro* experiments were sterilized with 100 % ethanol for 10 minutes under UV light exposure inside a biosafety cabinet. Then, the 3D microstructures were rinsed with sterile water to remove any traces of ethanol before cell seeding.

**Figure 1.**
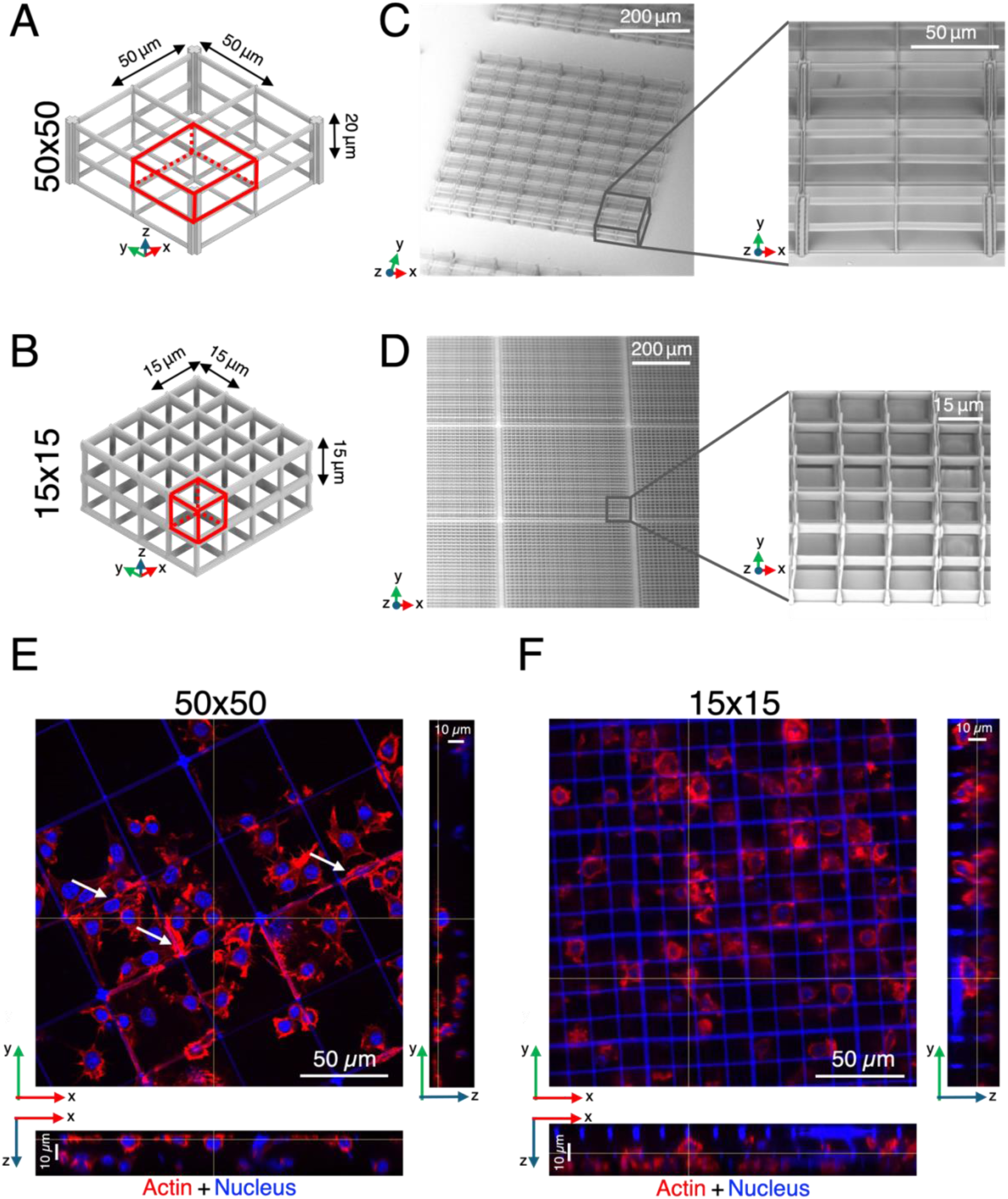
Design, fabrication, and macrophage colonization of 3D microstructures. (**A-B**) CAD models of the large pore (**A**) and small pore (**B**) microstructures. The elementary units are highlighted in red; each pore measures 50×50×20 μm³ (**A**) or 15×15×15 μm³ (**B**), respectively. (**C-D**). SEM images of the corresponding fabricated microstructures, demonstrating the fidelity of the two-photon polymerization process. (**E-F**) Confocal laser scanning microscopy images and orthogonal views of M0 macrophages cultured within the large pore (**E**) and small pore (**F**) scaffolds. Actin cytoskeleton is shown in red and nuclei in blue. In (**E**), white arrows indicate cells elongated along the beams of the large pore microstructure, while in (**F**) cells appear more spatially confined within the small pores.

### 2.2 Cell culture and staining of the cytoskeleton

Mouse macrophages RAW264.7 (TIB-71, ATCC) were maintained in T75 cell culture flasks (Biofil) and cultured in Dulbecco’s modified Eagle’s medium (DMEM, EuroClone, Italy), supplemented with 10 % FBS and 1 % penicillin/streptomycin (EuroClone, Italy) at 37 °C, 5 % CO_2_. Upon trypsinization (0.1 % trypsin/EDTA solution, EuroClone, Italy), cells were seeded on glass coverslips (Cover Slips, ThermoFisher Scientific, USA) with a diameter of 12 mm. Cells were plated on flat glass controls (Flat) and in the fabricated microstructures. A total amount of 20.000 cells per sample was plated in 500 μL complete medium. Before cell seeding, the substrates were sterilized with 100 % ethanol and UV irradiation for 10 minutes under a sterile biological hood and placed in a 24-well plate (EuroClone, Italy). All the samples were rinsed twice in PBS and fixed with paraformaldehyde solution (w/v 4 %) for 10 min; samples were then washed three times with glycine 0.1 M in PBS and permeabilized with 0.25 % Triton-X-100 in PBS for 10 min. Phalloidin-TRITC (P1951, dilution 1:100, Sigma-Aldrich, USA) was incubated for 1h at room temperature, to stain filamentous actin. After three rinses with 0.1 % Tween in PBS, cell nuclei were counterstained with 1 μM Hoechst 33342 (ThermoFisher Scientific, USA). Finally, samples were washed once in PBS and in distilled water (dH_2_O) and mounted with 10 μL MOWIOL DABCO. Confocal laser scanning microscopy images were acquired (Fluoview FV10i, Olympus, Japan) using a water immersion 60x objective (NA 1.2, 0.28 mmWD).

### 2.3 Chemical induction of macrophage polarization and analysis of cell phenotype

Upon cell seeding on the flat controls and on the fabricated microstructures, cells were let adhere overnight and then were chemically stimulated with 1 µg/mL Lipopolysaccharides from *Escherichia coli* O55:B5 (LPS, L6529, Sigma-Aldrich, USA) for promoting the pro-inflammatory M1 phenotype and with 40 ng/mL mouse interleukin-4 (IL-4, I1020, Sigma-Aldrich, USA) for inducing the anti-inflammatory M2 phenotype. After 24 hours of incubation at 37 °C, 5 % CO_2_, cell culture medium was replaced with fresh complete DMEM. Cultures were maintained in the incubator for further 48 hours and then analysed by immunofluorescence and FLIM microscopy to evaluate the effectiveness of cell polarization. Immunofluorescence assay was used for identifying the effect of 3D scaffolds and chemicals on cell polarization. All the samples (M0, M1 and M2 cells) were grown on all the investigated substrates: Flat, 15x15, and 50x50 scaffolds. Inducible nitric oxide synthase (iNOS) and Arginase 1 (Arg1) were used for the identification of M1 and M2 cell phenotypes, respectively. Moreover, actin staining was performed to investigate cell cytoskeletal organization. Cultures were rinsed twice in PBS and fixed with paraformaldehyde solution (w/v 4 %) for 10 min; samples were then washed three times with glycine 0.1 M in PBS and permeabilized with 0.25 % Triton-X-100 in PBS for 10 min. Blocking of not specific antibody binding sites was performed by incubating samples with a solution of 2 % bovine serum albumin (BSA) and 0.1 % Tween in PBS. Primary antibodies (Anti-iNOS [EPR16635] - ab178945-dilution 1:750, Abcam, UK; anti-Arg1 - ab96183 - dilution 1:750, Abcam, UK) were incubated overnight at 4°C. Upon rinsing three times in 0.1 % Tween in PBS, samples were incubated with secondary antibodies (Anti-rabbit Alexa Fluor^®^ 488-conjugated (ab150081, dilution 1:1000, Abcam, UK) in a solution of 2% BSA and 0.1 % Tween in PBS (45 minutes). After three more rinses with 0.1 % Tween in PBS, cell nuclei were counterstained with 1 μM Hoechst 33342 (ThermoFisher Scientific, USA). Finally, samples were washed once in PBS and in distilled water (dH_2_O) and mounted with 10 μL MOWIOL DABCO. Confocal laser scanning microscopy images were acquired on a microscope (Fluoview FV10i, Olympus, Japan) equipped with a water immersion 60x objective (NA 1.2, 0.28 mm WD). 405 and 473 nm diode lasers were used to excite Hoechst33342 and Alexa Fluor 488 secondary antibody. Emission DAPI (430-470 nm) and FITC (500-540 nm) filters were used for fluorescence detection. *Z*-stack images were collected with a *Z*-step of 1 μm, to acquire the samples throughout their full height. 1024x1024 pixel images were acquired with a resolution of 0.2 μm/pxl. Images were processed through ImageJ software (1.53, NIH, USA). Regions of interest (ROIs) were manually drawn along the cell boundaries, carefully adapted to each cell to exclude any fluorescence signal originating from SZ2080 autofluorescence. The fluorescence intensity of iNOS and Arg1 in the M0, M1, M2 macrophage samples were evaluated on images at the equatorial plane of cells in flat samples and close to the middle height of the pores in the two scaffolds. Fluorescence analysis was based on the quantification of the mean intensity value for each ROI, after subtraction of the background intensity. The values obtained for the M1 and M2 samples were then normalized with respect to the corresponding values of the M0 sample. The percentage induction efficiency was calculated by counting the number of fluorescent M1 and M2 macrophages and dividing it by the total number of cells present in each sample image, after subtracting the mean fluorescence intensity values of the corresponding M0 samples.

### 2.4 Cytoskeletal organization investigation and analysis

All the samples (M0, M1 and M2 cells grown on the different substrates, Flat, 15x15, and 50x50 scaffolds) were stained with Phalloidin-TRITC and Hoechst 33342, as previously described. Confocal laser scanning microscopy images were acquired (Fluoview FV10i, Olympus, Japan) using a water immersion 60x objective (NA 1.2, 0.28 mm WD). 553 nm diode lasers were used to excite the fluorescent dye and TRITC (555-620 nm) filter were used for the fluorescence detection. *Z*-stack images were collected with a *Z*-step of 1 μm, to acquire the samples throughout their full height microstructures to evaluate the localization of macrophages within the 3D volume. 1024x1024 pixel images were acquired with a resolution of 0.2 μm/pxl. Analyses were based on the detection of filamentous actin structure on images at the equatorial plane of cells both in Flat and in the middle height of the scaffolds. ImageJ software (1.53, NIH, USA) was used for image analysis. ROIs were manually drawn along the cell boundaries, carefully adapted to each cell to exclude any fluorescence signal originating from SZ2080 autofluorescence. For each selected ROI, cell area (area of selection) and two shape descriptors, circularity 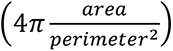 and maximum Feret’s diameter (longest distance between two parallel lines perpendicular to that distance and drawn at the object’s boundaries) were evaluated, as previously described [41, 42].

### 2.5 Cell metabolic activity investigation and analysis

RAW 264.7 murine macrophages in Flat, 15x15, and 50x50 scaffold were acquired using the two-photon fluorescence lifetime imaging (2P-FLIM) technique. Imaging was carried out 48 hours post induction of cell polarization by Nikon multiphoton microscope (Nikon, Japan) mounted on a Ti2-E inverted platform and equipped with a MultiHarp 150 time-correlated single photon counting (TCSPC) module and a hybrid PMA detector (PicoQuant, Germany). Endogenous NAD(P)H fluorescence was excited using a mode-locked Ti:Sapphire laser (Chameleon Vision II, Coherent, USA) tuned to 750 nm and operating at 80 MHz. The laser beam was focused through a 40x water immersion objective (NA 1.15, WD 0.62 mm, Nikon). Emitted fluorescence was filtered using a 440/40 nm bandpass filter, enabling selective detection of NAD(P)H emission in the blue region of the visible spectrum. Cells were maintained at 37°C in a humidified chamber (Okolab, Italy) with 5 % CO_2_ and 95 % relative humidity throughout image acquisition. FLIM image stacks (512x512 pixels) were acquired over a 10-minute time window, with a pixel dwell time of 27.2 µs and 0.621 µm/pixel. A 2x binning was further applied prior to analysis to maximize the photon count per pixel. Five ROIs per image were manually drawn to perform single-cell analysis and to exclude fluorescence artifacts arising from the microstructure’s photoresist. Each ROI typically reached approximately 10^3^ photons/pxl. Since the scaffold exhibited significantly higher fluorescence intensity compared to endogenous fluorophores, such as NAD(P)H, additional ROIs were manually and intensity-based defined on the scaffold beams. Their fluorescence lifetime was measured separately to account for potential signal leakage from the scaffold into the cellular ROIs, thereby preserving the accuracy of the metabolic activity analysis. Fluorescence decay curves were analysed using SymPhoTime 64 software (v2.7, PicoQuant), which applies an iterative reconvolution algorithm to fit the experimental photon arrival histograms. The distribution of the excited state lifetimes, *I(t)*, was modelled according to a double-exponential decay function including the instrument response function (IRF):

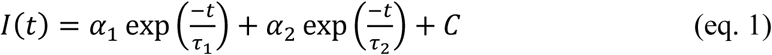

*α*_1_ and *α*_2_ represent the relative amplitudes of short and long lifetime components, respectively. *τ*_1_ and *τ*_2_ are the short and long lifetime components, respectively; *C* corresponds to the background. The biexponential model was employed to account for the two main species of NAD(P)H: a short lifetime component (*τ₁*, typically ∼0.4 ns), corresponding to free NAD(P)H in the cytosol, and a long lifetime component (*τ₂*, typically ∼2.0-3.0 ns), associated with enzyme-bound NAD(P)H. To characterize the metabolic activity of cells the mean fluorescence lifetime was estimated by means of the amplitude average [43] using:

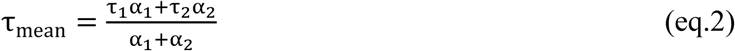

As the lifetime of free NAD(P)H does not change by more than 0.2 ns across the different samples, it was fixed at 0.42 ns (average value across the samples) This helped in improving the accuracy of protein-bound NADH lifetime estimation, leading to more reliable quantification of metabolic shifts by constraining one component of the bi-exponential decay model. The goodness of fit was examined using a chi-squared statistical test; a χ^2^ <1.2 was deemed to be a satisfactory fit (see **Supplementary** Figure 1).

### 2.6 Phasor Plots

Additionally, FLIM data was analysed using phasor plots, a graphical and fit free method to visualize FLIM data. In essence, phasor plots are created by calculating a Fourier transform of decay data in the time domain acquired image. For each pixel, the phasor coordinates which are given by (*g*, *s*) were calculated using:

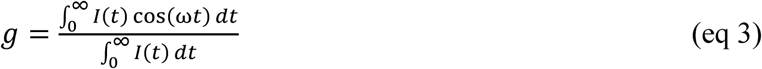

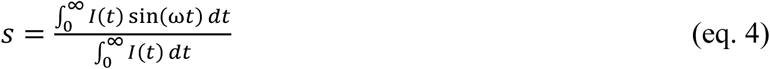

where ω = 2π𝑓 is the angular frequency of the laser repetition rate. These coordinates measured on individual pixels were the plotted on a Fourier space {*g, x*}. Fluorophores with mono exponential decay lie on a universal semicircle in this Fourier space, while multiexponential decays, such as a mixture of free and bound NADH proteins, lie on a cord joining points on the semicircle. This fit-free approach is faster, more robust against noise and offers easy interpretability [44, 45]. Taking advantage of the monoexponentially decays lying on the universal circle, a dye characterized by a well-known mono exponential decay can be used to calibrate the phasor plot. For calibration, a 10 µM fluorescein solution diluted in deionized water at a final pH 9 was used. The analysis was performed using the open-source software FLUTE [46]. It allows to create circular ROIs with a fixed radius on the phasor to eliminate datasets coming out from the fluorescence of the scaffolds. This proved to be important to show clean and reliable results on cell metabolic activity. Additionally, an in-house written script was created on top of FLUTE’s graphical user interface, to visualize multiple phasor plots on one for easier comparison between different samples. This allows for probing the metabolism of the cells in the three different conditions (Flat, 15x15 and 50x50). After loading images on custom FLUTE, a 3x3 median filter was applied iteratively four times to all images acquired to reduce noise. Subsequently, intensity-based thresholding was applied to filter out low-intensity pixels. The minimum intensity thresholds used were 50 counts for Flat, 120 counts for the 15x15 scaffold, and 150 counts for the 50x50 scaffold images. To remove scaffold-related signals in the phasor plot analysis, a distance threshold was applied. Specifically, a distance of 0.72 μm was used for the 50x50 scaffold and 0.73 μm for the 15x15 scaffold, effectively excluding pixels associated with scaffold autofluorescence.

### 2.7 Statistical analysis

The statistical analysis of the collected data was performed with OriginPro (OriginPro 2024b software, OriginLab Corporation, Northampton, Massachusetts, USA) applying non parametric tests with Kruskal-Wallis ANOVA with Dunn’s multiple comparison test for (data not normally distributed) and ordinary one-way ANOVA with Tukey’s multiple comparisons test (data normally distributed), with the following significance: \**p*-value < 0.05; \*\**p*-value < 0.01; \*\*\**p*-value < 0.001; \*\*\*\**p*-value < 0.0001. Each type of experiment (cell polarization, cytoskeletal organization, cell metabolic activity) was performed at least in triplicate. More than 85, 419, and 34 cells were quantified for cell polarization, cytoskeletal organization, and metabolic activity experiments, respectively, for each combination of cell population and substrate. To better highlight differences between samples in the cell polarization and cytoskeletal organization experiments, we calculated the percentage variation (%), defined as the median ratio of the numerical values obtained from the respective quantifications.

## 3. Results and discussion

### 3.1 Fabrication of 3D microstructures and colonization by macrophages

3D scaffolds composed by two different pore sizes (50x50x20μm^3^ is shown in **Figure 1A**, and 15x15x20μm^3^ in **Figure 1B**) were fabricated on the top of circular glass coverslips by two-photon laser polymerization. After sample development, the microstructures were inspected by scanning electron microscopy (**Figures 1C, 1D**), confirming that they matched the designed specifications in terms of pore dimension and structural stability. RAW264.7 macrophages were grown in the 3D samples (namely, 50x50 and 15x15). 72 hours after seeding cell nucleus and actin cytoskeleton were labelled with fluorescent dyes and confocal laser scanning microscopy was performed for evaluating scaffold colonization and spatial distribution of cells. Cells efficiently occupied the 3D volume of both the microstructures, as can be observed in the 3D reconstructions of the acquired *Z*-stacks (**Supplementary Video 1, 2**) and in the orthogonal projections (*XZ* and *YZ* images of the acquired confocal laser scanning microscopy images, **Figures 1E, 1F**). Moreover, cell morphology differences were also evident from the images. In the 50x50 scaffold, some cells tended to elongate along the beams of the microstructure whenever space allowed (white arrows), whereas in the 15x15 scaffold, individual cells appeared confined within single pores.

### 3.2 3D microstructures induce morphological changes of the actin cytoskeleton and metabolic activity in M0 macrophages

To characterize the influence of the 3D geometry on macrophage behaviour, we investigated cell morphology [47] and metabolism [27, 48]. In recent years, the use of FLIM to investigate macrophage polarization has become an increasingly prominent topic, attracting growing attention within the scientific community; a steadily expanding body of literature demonstrates how this technique can be leveraged to distinguish between pro-inflammatory and anti-inflammatory phenotypes [49–52]. Notably, machine learning approaches have been employed to train algorithms capable of identifying macrophage polarization states in real time, further highlighting the potential of FLIM as a powerful, non-invasive tool for immune cell characterization [48, 51]. This work is the first to our knowledge that completely avoids considering the contribution of the flat surfaces surrounding the fabricated 3D scaffolds, by analysing only macrophages effectively located inside the microstructures by means of imaging techniques. Firstly, we analysed the actin cytoskeleton organization in M0 cells seeded in the three substrates (Flat, 50x50 and 15x15). Confocal laser scanning microscopy imaging of samples stained with phalloidin-TRITC was performed, to evaluate the organization of the actin cytoskeleton. Qualitatively, M0 macrophages appeared mainly as rounded cells with few short pseudopods and rare membrane extensions in the flat control, while in the 50x50 and 15x15 microstructures they displayed a few elongations and some membrane extrusions, especially in the microstructures characterized by larger pores, developed to attach to the scaffolds’ beams (**Figure 2A**). Quantitative analysis of cell area showed a significant enlargement in both 3D microstructured environments respect to the flat substrate, with the effect being more marked in the small pore configuration. This effect likely reflects different modes of cell-scaffold interaction: in scaffolds with small pores, cells occupy the entire available space, whereas in scaffolds with large pores, they elongate along the beams, when spatially permitted. These distinct morphological adaptations suggest that the 3D geometry can induce a remodelling of the cytoskeleton adapting to the size of the pore (**Figure 2B**, and **Supplementary** Figure 2A). Therefore, cell morphology was further characterized by evaluating macrophage circularity and maximum Feret’s diameter. Circularity estimates the degree of roundness of the cells, with values of 1 indicating a perfect circle and values approaching 0 corresponding to increasingly elongated shape. Macrophages cultured within the two 3D microstructures exhibited a significantly more elongated morphology compared to the flat control, as shown by a percent reduction in circularity of 8 % for 50×50 and 18 % for 15×15 pores. This was most pronounced in the 15x15 pores (**Figure 2C**, and **Supplementary** Figure 2B) suggesting substantial cytoskeleton rearrangement due to cell confinement within the small pore space. Furthermore, we obtained similar results analysing the maximum Feret’s diameter, which is a measurement of the longest distance between any two points along the selected boundary and that it is expected to increase proportionally to cell elongation [42, 47]. In our experiments, both 3D microstructures induced an increase in the maximum Feret’s diameter, without any significant differences between the two pore sizes (**Figure 2D**, and **Supplementary** Figure 2C). This result further corroborated the influence of the 3D geometry inducing modifications at the level of the cytoskeleton in M0 macrophages. These morphological changes may impact both cellular metabolic activity and polarization. Therefore, as a next step, we investigated cellular metabolic activity using FLIM microscopy. In recent years, the use of FLIM to investigate macrophage polarization has become an increasingly prominent topic, attracting growing attention within the scientific community; a steadily expanding body of literature demonstrates how this technique can be leveraged to distinguish between pro-inflammatory and anti-inflammatory phenotypes [49–52]. Notably, machine learning approaches have been employed to train algorithms capable of identifying macrophage polarization states in real time, further highlighting the potential of FLIM as a powerful, non-invasive tool for immune cell characterization [48, 51]. This work is the first to our knowledge that completely avoids considering the contribution of the flat surfaces surrounding the fabricated 3D scaffolds, by analysing only macrophages effectively located inside the microstructures by means of imaging techniques. Our results revealed that M0 macrophages cultured on both 50x50 and 15x15 scaffolds exhibited (≃10 %) shorter mean fluorescence lifetimes compared to those on flat control, suggesting a shift toward increased glycolytic activity (**Figure 2E**). This effect appeared to be related solely to the dimensionality of the cellular environment (3D for the photopolymerized scaffolds *versus* 2D for the flat substrate). Indeed, the overall metabolic profiles of M0 macrophages were similar across both scaffold substrates, indicating that pore size has minimal impact on the metabolic activity of unpolarized macrophages.

**Figure 2.**
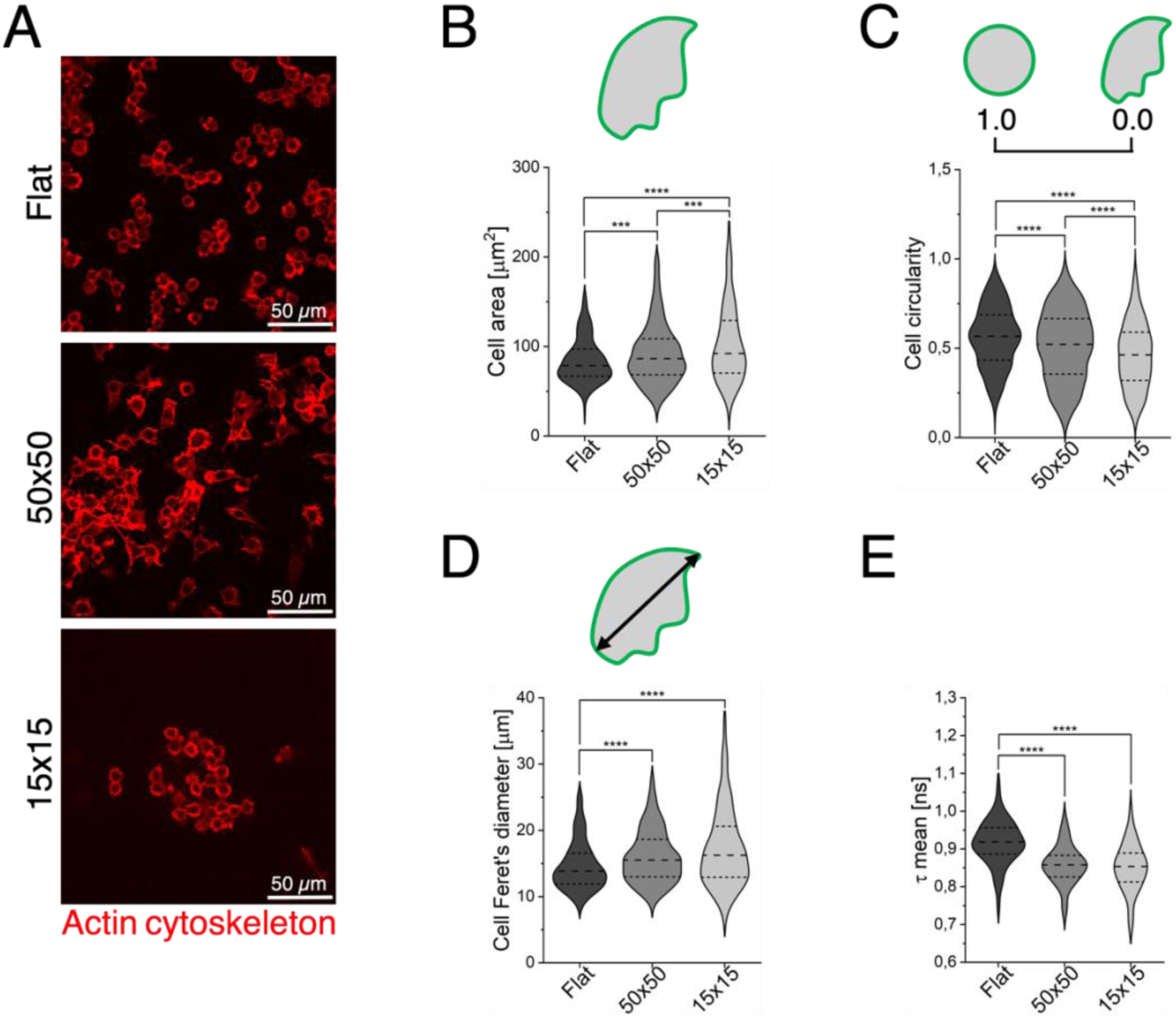
Morphology and metabolic activity of M0 macrophages in the presence of the 3D microstructures. **A**) Representative confocal laser scanning microscopy images of M0 macrophages cultured on Flat, 50x50, and 15x15 substrates, stained with Phalloidin-TRITC to visualize the actin cytoskeleton. **B-D**) Quantification of M0 macrophages morphological parameters: cell area (**B**), circularity (**C**), and maximum Feret’s diameter (**D**). **E**) Analysis of microstructure-induced modulation of cellular metabolism in M0 macrophages.

### 3.3 3D microstructures effects on macrophage polarization

From the results obtained, we inferred that the 3D microstructures demonstrated to be able to induce alterations in the morphology and metabolism of M0 macrophages. We then asked whether microstructures *per se* could induce any spontaneous polarization. Therefore, to detect the eventual polarization phenotypes of macrophages, the basal expression of two biomarkers was assessed by immunostaining with antibodies specific for inducible nitric oxide synthase (iNOS), identifying M1 macrophages, and arginase 1 (Arg1), detecting M2 cells. Representative images of the obtained iNOS and Arg1 fluorescence signals in M0 cells are reported in **Figure 3A**. The fluorescence intensity was quantified from the images acquired by confocal laser scanning microscopy of the three assayed substrates. Only cells inside the microstructures were considered (see **Materials and Methods section 2.3**). This analysis allowed to determine whether the 3D microstructures can induce any spontaneous polarization of the cultured macrophages. As shown in **Figures 3B-3C**, fluorescence intensity quantification revealed that neither the 50x50 nor the 15x15 scaffolds significantly altered neither iNOS nor Arg1 expression levels, suggesting that the 3D microstructures alone do not induce spontaneous phenotypic changes. These results demonstrated that our 3D microstructures not only can be used as an “inert” model for studying *in vitro* the interaction with macrophages and other cell types, but they can also be involved, for example, in the tissue regeneration (*i.e.,* endothelial cells and fibroblasts) or the pathology fields (*i.e.,* cancer). Having characterized the baseline effects of our scaffolds on macrophage behaviour, we made the model more complex by introducing biochemical cues, which are indeed present in realistic and physiologically relevant conditions. Therefore, M0 cells were chemically stimulated with lipopolysaccharides (LPS) for inducing their polarization towards the M1 pro-inflammatory phenotype, and interleukin-4 (IL-4) for polarizing them towards the anti-inflammatory M2 phenotype. The efficiency of the chemical stimulation protocol, in combination with the three different substrate types, was then evaluated. Representative images acquired by confocal laser scanning microscopy are reported in **Figure 3D**. For each induced phenotype, the percentage of polarized macrophages related to each tested substrate was calculated based on the number of fluorescent cells (see **Materials and Methods section 2.3**). As shown in **Figures 3E-3F**, upon stimulation with either LPS or IL-4, significant differences in induction efficiency were observed between the 3D microstructures and the Flat, demonstrating that both induction processes were markedly enhanced under 3D conditions. Specifically, both microstructures amplified the cytokine effects in promoting macrophage polarization: LPS induction increased by 26 % in 50×50 and by 40 % in 15×15 compared to Flat; IL-4 induction increased by 18 % in 50×50 and by 30 % in 15×15 *versus* Flat.

**Figure 3.**
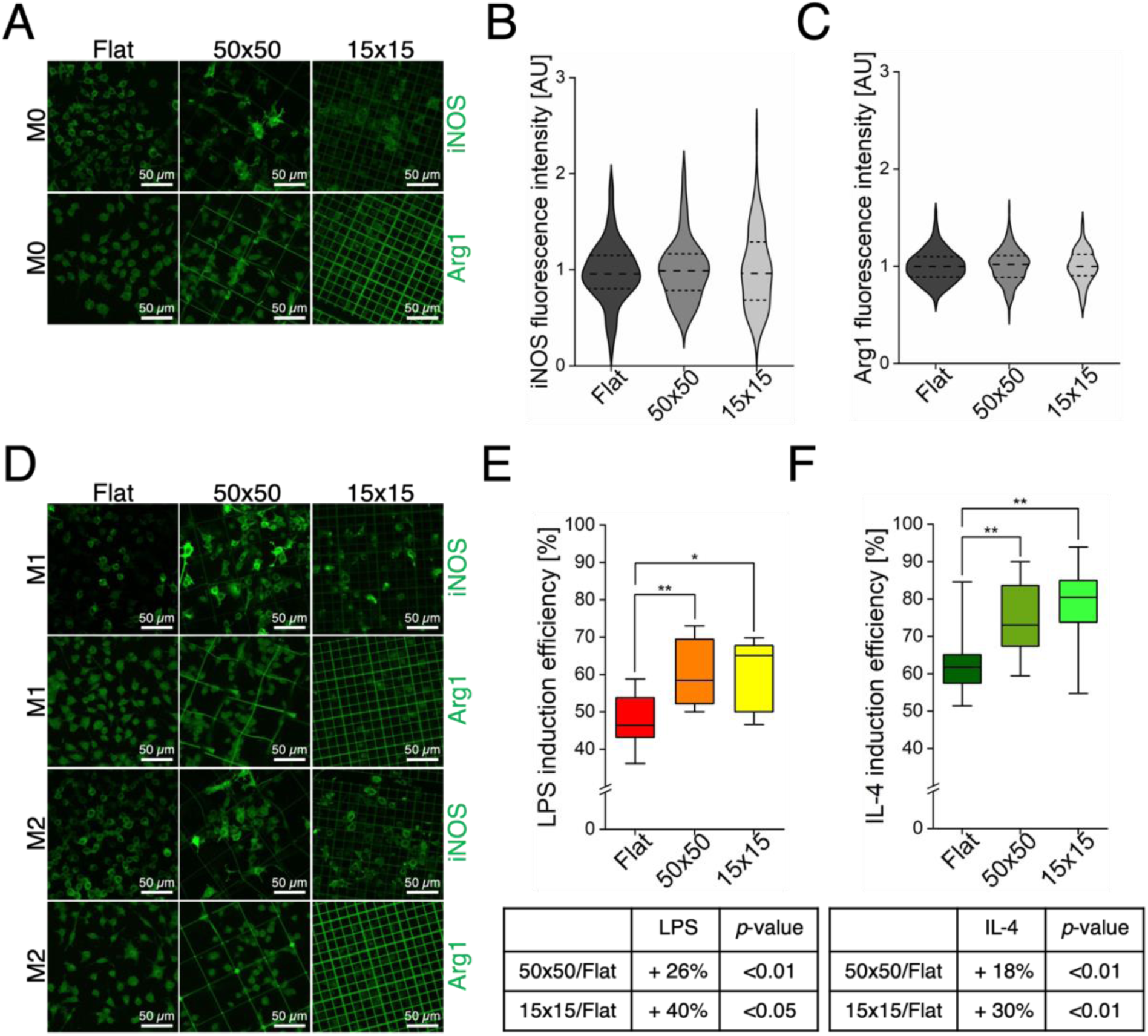
3D microstructures do not polarize M0 macrophages. **A**) Representative confocal laser scanning microscopy images of M0 macrophages cultured on Flat, 50x50, and 15x15 substrates, stained with antibodies against iNOS and Arg1. **B-C**) Quantification of fluorescence intensity for iNOS (**B**) and Arg1 (**C**), normalized to Flat control, in M0 macrophages cultured on Flat and within the 50x50 and 15x15 microstructures. **D**) Representative confocal laser scanning microscopy images of M1 and M2 macrophages cultured on the same substrates and stained for iNOS and Arg1. **E-F**) Quantification of induction efficiency (% of green polarized cells/total cells) following LPS (**E**) and IL-4 (**F**) treatment on Flat, 50x50, and 15x15 substrates, based on anti-iNOS and anti-Arg1 staining.

### 3.4 3D microstructures induce morphological and metabolic changes and efficiently modulate macrophage phenotype in combination with chemical stimulation

In recent years, considerable efforts have been made to understand how morphometric analysis, either alone or in combination with the expression of specific markers, can provide direct insights into macrophage polarization. For instance, Rostam *et al.* developed machine learning algorithms capable of automatically identifying functional phenotypes based on cell size and morphology [53], while Geng *et al.* employed a polymer-protein sensor array to retrieve unambiguous information on macrophage polarization states [54]. More recently, Dannhauser *et al.* developed a lab-on-a-chip approach based on single-cell, label-free and high throughput light scattering pattern analyses to identify macrophages, combined with a machine learning-based approach to efficiently classify their phenotypes [55]. However, the relevance of these approaches may be limited if the morphological rearrangement of cells within three-dimensional, *in vivo*-like environments is not considered. To address this, we chose to perform experiments that combine both chemical stimulation and physical cues provided by engineered 3D microenvironments, aiming to more closely mimic the physiological conditions encountered following biomaterial implantation. In this context, we take a step further by systematically investigating, in a controlled *in vitro* setting, how defined scaffold architectures shape macrophage morphology and activation. M0 cells were chemically stimulated with LPS and interleukin 4. As can be observed in **Figure 4A**, M1 cells on the flat substrate showed membrane extensions and elongated pseudopods, adopting a larger and more flattened morphology compared to the M0 cells. M1 macrophages grown in the 50x50 microstructures displayed a similar morphology and an irregular shape with membranes extending along the microstructure’s beams. On the contrary, in the case of the 15x15 scaffolds, cells displayed limited morphological modifications compared to the M0 cells, maintaining a roundish morphology due to the constrictions given by the small pore size. In the case of M2 polarization, cells exhibited increased membrane protrusions and elongations, appearing more flattened and fusiform (spindle shaped) both on the flat control and in the 50x50 scaffold, while they displayed limited membrane extensions in the 15x15 scaffold. To support the qualitative observations, we quantitatively analysed macrophage morphology using area, circularity, and maximum Feret’s diameter, the same parameters considered for M0 characterization in the previous section. All the comparisons between the induced samples in terms of variation (%) were calculated for all the three measured parameters and are reported in **Supplementary** Figures 2A**, 2B, 2C**. M1 macrophages showed a significant increase in cell area compared to not stimulated M0 cells, with variations of +15 % on Flat and +14 % in the 50x50 microstructure, while no significant change was observed in the 15x15 scaffold (**Figure 4B**, left). Similarly, compared to not stimulated M0 cells, M2 macrophages exhibited a significant increase in area only on Flat and 50x50 substrates (+17 % and +12 %, respectively), with no appreciable variation in the 15x15 condition (**Figure 4B**, right). On the contrary, investigating cell circularity, we observed that M1 cells (**Figure 4C**, left), displayed a significant increase (+ 20 %) in 15x15 microstructures compared to the not stimulated M0 cells, thus suggesting that the chemical stimulus, combined with the small pore geometry, is the only condition where we can observe a pro-inflammatory phenotype. Additionally, a circularity decrease with respect to the M0 phenotype was detected in M2 cells both on the flat substrate and in the 50x50 scaffold, -8 % and -22 %, respectively, (**Figure 4C**, right): this behaviour was not detectable in the 15x15 microstructures. The maximum Feret’s diameters measured of pro-inflammatory macrophages (**Figure 4D**, left) exhibited a statistically significant increase only on the flat condition (+7 % compared to M0), thus suggesting that the enhancement in cell elongation is only partially driven by chemical stimulation. Concerning the anti-inflammatory (M2) phenotype, the highest maximum Feret’s diameters were found in both Flat and 50×50 conditions (**Figure 4D**, right), with increases of +7 % and +9 %, respectively, indicating a modest but significant synergy between chemical and physical cues. From these results, it can be inferred that the amount of free volume available for the cells in the microstructures is clearly reduced within the 15x15 pores, creating a more confined and stressful environment and promoting a pro-inflammatory M1 phenotype. To investigate how environmental modulation affects macrophage metabolism, we employed FLIM as a label-free approach to assess the metabolic state of the cells as shown in **Figure 4E**, a reduction in the average fluorescence lifetime was observed not only in non-polarized macrophages (M0), but also in pro-inflammatory (M1) and anti-inflammatory (M2) phenotypes cultured within both microstructured scaffolds, when compared to the flat substrate. These results confirm that the three-dimensionality of the cellular microenvironment significantly impacts cell metabolism, promoting a shift towards a more glycolytic state. To ensure that the scaffold material itself was not contributing to the observed differences in cellular lifetimes, fluorescence lifetime measurements of the two scaffolds were also performed. As shown in **Supplementary** Figure 2 **E and 2F**, the 15x15 scaffold exhibited a higher mean lifetime compared to the 50x50 one. This result is not unexpected and can be attributed to the different parameters used during the photopolymerization process of the scaffolds, such as laser power, scan speed, and printing geometry. However, what is crucial to underline is that our cell metabolic measurements are not affected by artifacts arising from scaffold autofluorescence. In the case of the 15x15 microstructure, for instance, accurately identifying cell boundaries is more challenging due to the proximity of the beams, making it more difficult to exclude signal contributions from the used material. Nevertheless, as shown in **Supplementary** Figure 2F, the fluorescence lifetime of the scaffold in the small pore microstructure (15x15) is significantly higher than the metabolic signal derived from the cells, thus confirming that, in the selected ROI, the cellular lifetime values are not compromised by scaffold interference. For the large pore microstructures (50x50), manual ROI selection was considerably easier, and therefore less concerning. However, since the fluorescence lifetime of the material, in this case, is similar to that of the cells, we further validated our data using an alternative approach. As shown in **Supplementary** Figure 3, a cell-scaffold thresholding procedure was also performed using the FLUTE software, based on phasor analysis. The resulting images clearly distinguish the cellular component (in green) from the polymeric scaffold (in red), allowing for the exclusion of the scaffold contribution. This confirms the reliability and robustness of the obtained metabolic data. This observation warranted further analysis to confirm that the variations seen in cellular data were primarily due to biological differences rather than possible scaffold autofluorescence leaking into the ROIs. Given the strength of the phasor approach, we applied it to evaluate whether there are differences in metabolic activity between cells cultured in the two microstructured scaffolds. This fit-free analysis allowed for intuitive classification of signal sources by applying circular ROIs directly onto the phasor space, as reported in **Supplementary** Figure 3. After classifying the cells and scaffold regions, a distance-based filtering method was applied to exclude scaffold signals, ensuring that subsequent analyses focused solely on cellular fluorescence lifetimes. Finally, these phasor clouds were plotted on a single phasor plot (**Supplementary** Figure 2 **D**), to compare the metabolic activity of macrophages grown on Flat, 50x50, and 15x15 samples. For each dataset, the centroid of the phasor cloud was calculated and marked with a cross on the phasor plot. It is equivalent to the mean lifetime calculated with the previous approach and serves as a reference point for comparing the cells grown in different conditions. A shift of the phasor plot centroid towards shorter lifetime values indicates a metabolic transition of the cells toward a more glycolytic state. This figure clearly shows that, for each macrophage phenotype (M0, M1, and M2), the 3D microenvironment induced a shift toward a more glycolytic metabolic state compared to flat control. Notably, the small pore scaffold further enhanced this glycolytic activity with respect to the large pore scaffold. The increased glycolytic activity observed in macrophages within the 15x15 scaffolds suggested a metabolic environment less supportive of M2 polarization, which typically favours oxidative phosphorylation.

**Figure 4.**
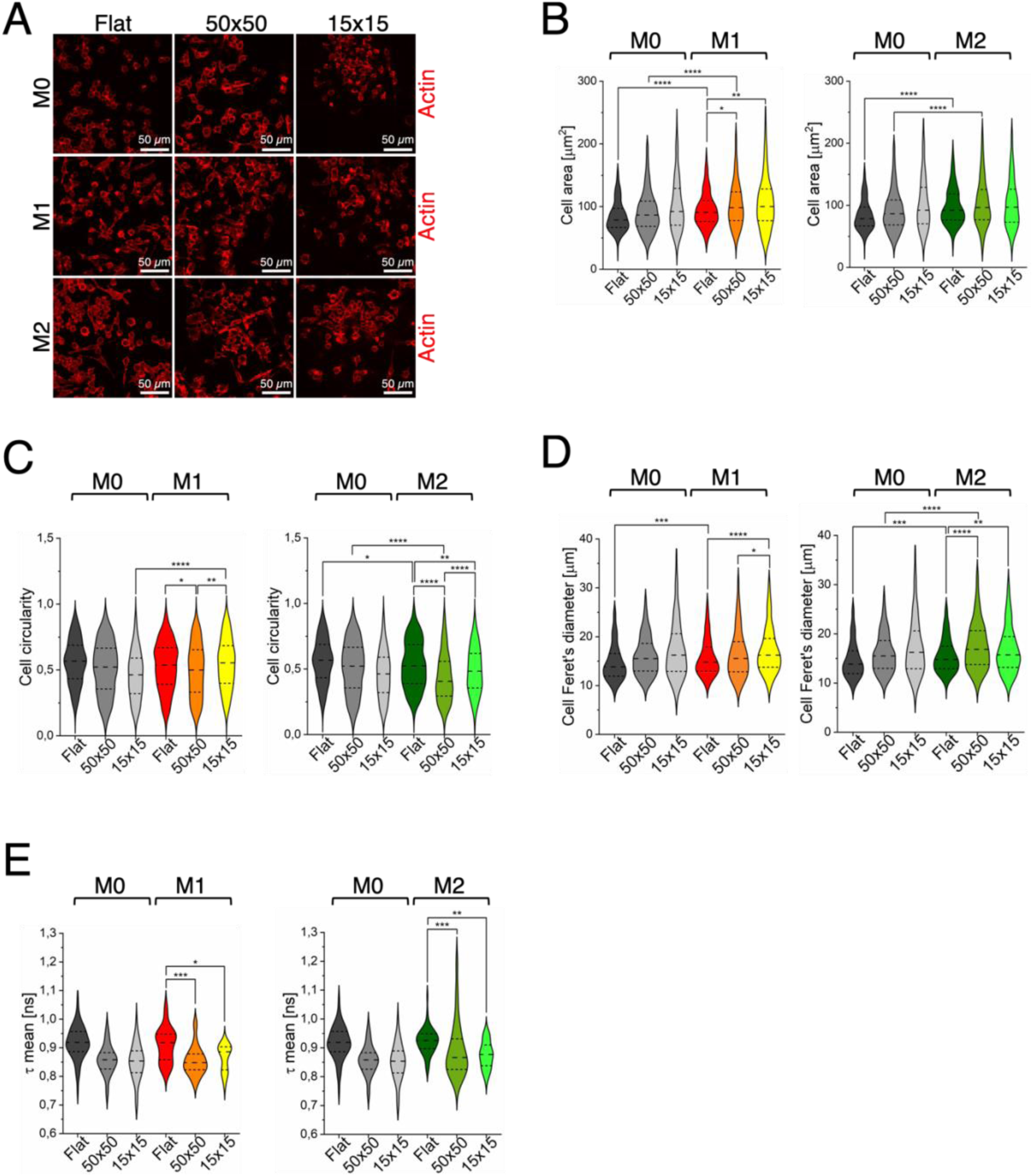
3D microstructures modify morphology and metabolism of macrophages in combination with chemical stimulation. **A**) Representative confocal laser scanning microscopy images of M0, M1, and M2 macrophages cultured on Flat, 50x50, and 15x15 microstructures, and stained with Phalloidin-TRITC to visualize the actin cytoskeleton. **B-D**) Quantification of morphological parameters in M0, M1, and M2 macrophages: cell area (**B**), cell circularity (**C**), and maximum Feret’s diameter (**D**), across the three substrates. **E**) Quantification of the metabolic activity in M0, M1, and M2 macrophages cultured in the same substrate conditions.

The combined effect of chemical stimulation and 3D architecture on macrophage polarization was then assessed by quantifying iNOS expression in LPS induced M1 macrophages and Arg1 expression in IL-4 treated M2 macrophages (**Figure 5A, 5B**). In the case of LPS treatment, data suggested that the 50x50 scaffold partially attenuated the effect of the chemical stimulus (**Figure 5C**). This was evident from the quantification of iNOS fluorescence intensity, which increased in all the tested conditions, but to a lesser extent in the 50x50 substrate compared to the flat control and the 15x15 scaffold (+119 % in Flat *vs.* +80 % in 50x50 and +120 % in 15x15) (**Figure 5D**). Striking results were obtained when analysing cells treated with IL-4 to induce an anti-inflammatory phenotype. As shown in **Figures 5E and 5F**, while both the Flat and the 50x50 supported a robust and comparable Arg1 expression response (+29 % and +27 % respect to M0 cells, respectively), the 15x15 microstructure failed to elicit any detectable upregulation of Arg1. This suggested that the small pore size might impair the effectiveness of M2 polarization, prevailing over the chemical stimulus.

**Figure 5.**
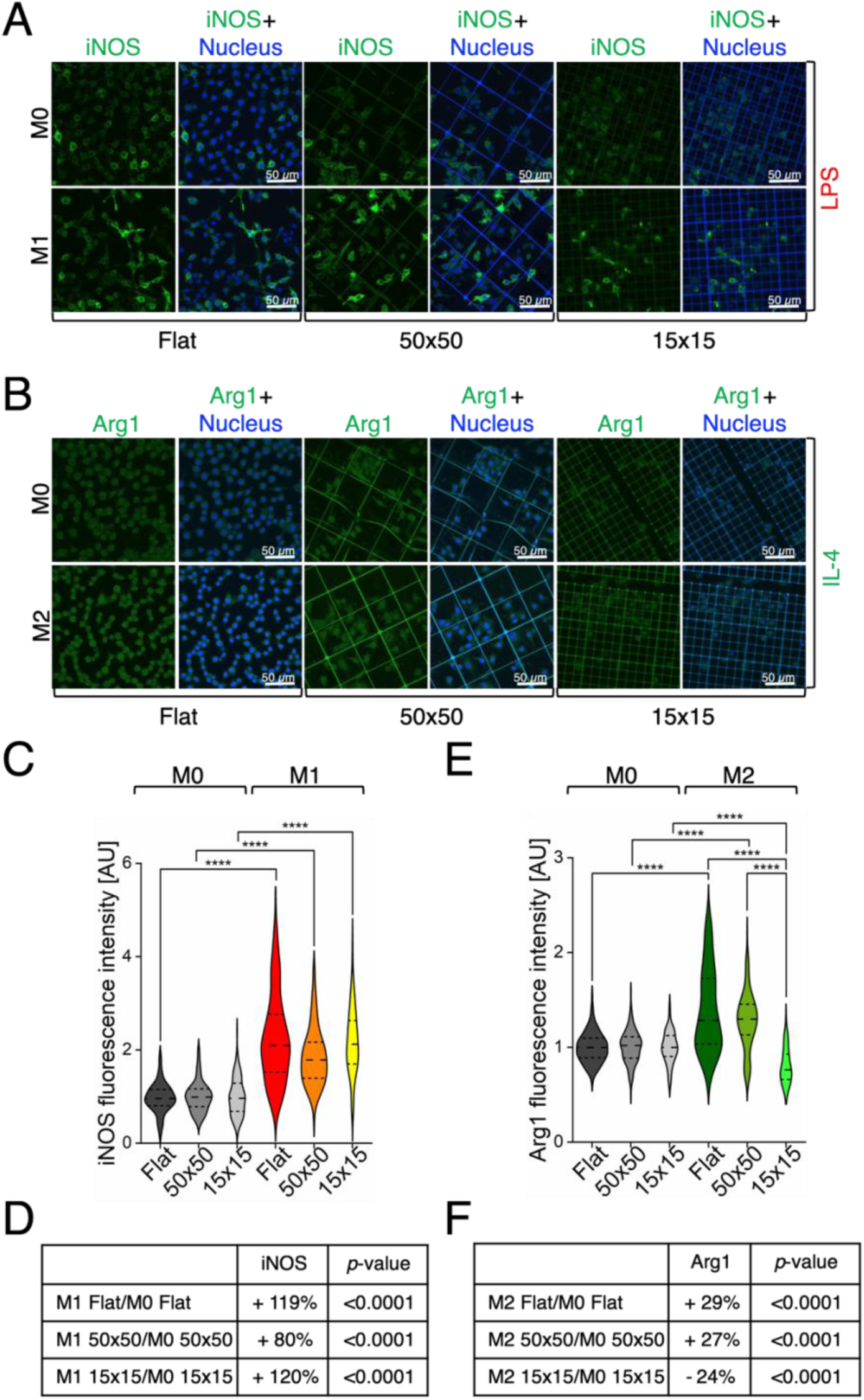
**3D microstructures modulate macrophage phenotype in combination with chemical stimulation. A-B**) Representative confocal laser scanning microscopy images of M0, M1, and M2 macrophages cultured on Flat, 50x50, and 15x15 samples and stained for iNOS (green, **A**) or Arg1 (green, **B**) along with nuclear counterstaining (blue), following chemical stimulation with LPS (M1) or IL-4 (M2). **C-D**) Quantification of iNOS fluorescence intensity in M1 macrophages cultured on the three samples investigated, normalized to the corresponding M0 controls (**C**). The variations (%) of iNOS expression are summarized in table (**D**). **E-F)** Quantification of Arg1 fluorescence intensity in M2 macrophages cultured on the three samples investigated, normalized to the corresponding M0 controls (**E**). The variations (%) of Arg1 expression are summarized in the table (**F**).

### 3.5 3D microstructures with large pores efficiently modulate macrophage phenotype towards an anti-inflammatory phenotype in combination with chemical stimulation

A further analysis was conducted to investigate the two biomarkers’ expression upon administration of the opposite stimulus, meaning the expression levels of Arg1 upon LPS stimulation and of iNOS upon IL-4 stimulation. To our knowledge, this is the first “cross” analysis of macrophage specific biomarkers performed with the goal to unravel the influence of 3D geometries combined with chemical induction. The expression levels of Arg1 (**Figure 6A**) and iNOS (**Figure 6B**) were evaluated in all the three substrates (Flat, 50x50 and 15x15) and were compared to the respective levels measured in M0 cells. Results of Arg1 fluorescence intensity quantification, measured in M1-stimulated cells, are reported in **Figure 6C** and **6D**, revealing remarkable findings. Indeed, a slight significant increase in Arg1 expression, compared to untreated cells, was observed only in the 50x50 microstructures (+5 %, **Figure 6D**). This result indicates that, within a complex system combining both physical and chemical stimuli as found *in vivo*, only the 50x50 microstructure significantly influences macrophage behaviour promoting an M2 anti-inflammatory polarization. Conversely, the smaller pore architecture favours a massive switch towards a pro-inflammatory response, as evidenced by elevated iNOS expression in IL-4–induced M2 macrophages, further highlighting the pivotal role of pore size in directing macrophage phenotype. Results reported in **Figure 6E** and **6F**, show that only in the 3D microenvironment there was a significant increase of the iNOS expression level with respect to the untreated cells, even more strikingly in the microstructure with small pores (+33 % in 50x50 and +121 % in 15x15), thus suggesting that the small pores strongly support a pro-inflammatory phenotype. Moreover, a statistically significant difference of iNOS expression levels can be appreciated between both the microstructured substrates with respect to the flat condition, thus indicating a synergistic role of the two stimuli. Summarizing, these data suggest that macrophages, upon chemical stimulation, are prompted to polarize toward an anti-inflammatory phenotype by physical interaction with the 50x50 scaffold, and toward an inflammatory phenotype by interaction with the 15x15 microstructure. Our findings are consistent with previous studies performed on porous polymeric scaffolds, reporting that pore size plays a crucial role in directing macrophage polarization, with large pores generally favouring the M2 anti-inflammatory phenotype, likely due to increased system permeability [21, 22]. Here we show that our ability to precisely design and control the 3D scaffold architecture can be exploited to modulate macrophage responses. In particular, the dimensions of the microstructure’s beams and the ability to fabricate high resolution geometries appear to be key parameters for promoting cell adhesion and regulating cellular activity. Notably, this is the first work describing the use of microstructures with defined pore sizes providing physical cues to macrophages in an environment characterized by the co-existence of chemical stimulation and thus mimicking the physiological reactions that take place when a biomaterial is implanted *in vivo*. The possibility of precisely tuning down at the micro- and submicrometric scales the pore dimension of the microstructures, thanks to two-photon polymerization, may allow to fabricate 3D scaffolds for decorating medical devices to be implanted and to finely modulate immune responses to favour a successful integration into the host tissues. Moreover, it provides a potential innovative strategy for the development of reliable *in vitro* platforms for anti-inflammatory drug screenings.

**Figure 6.**
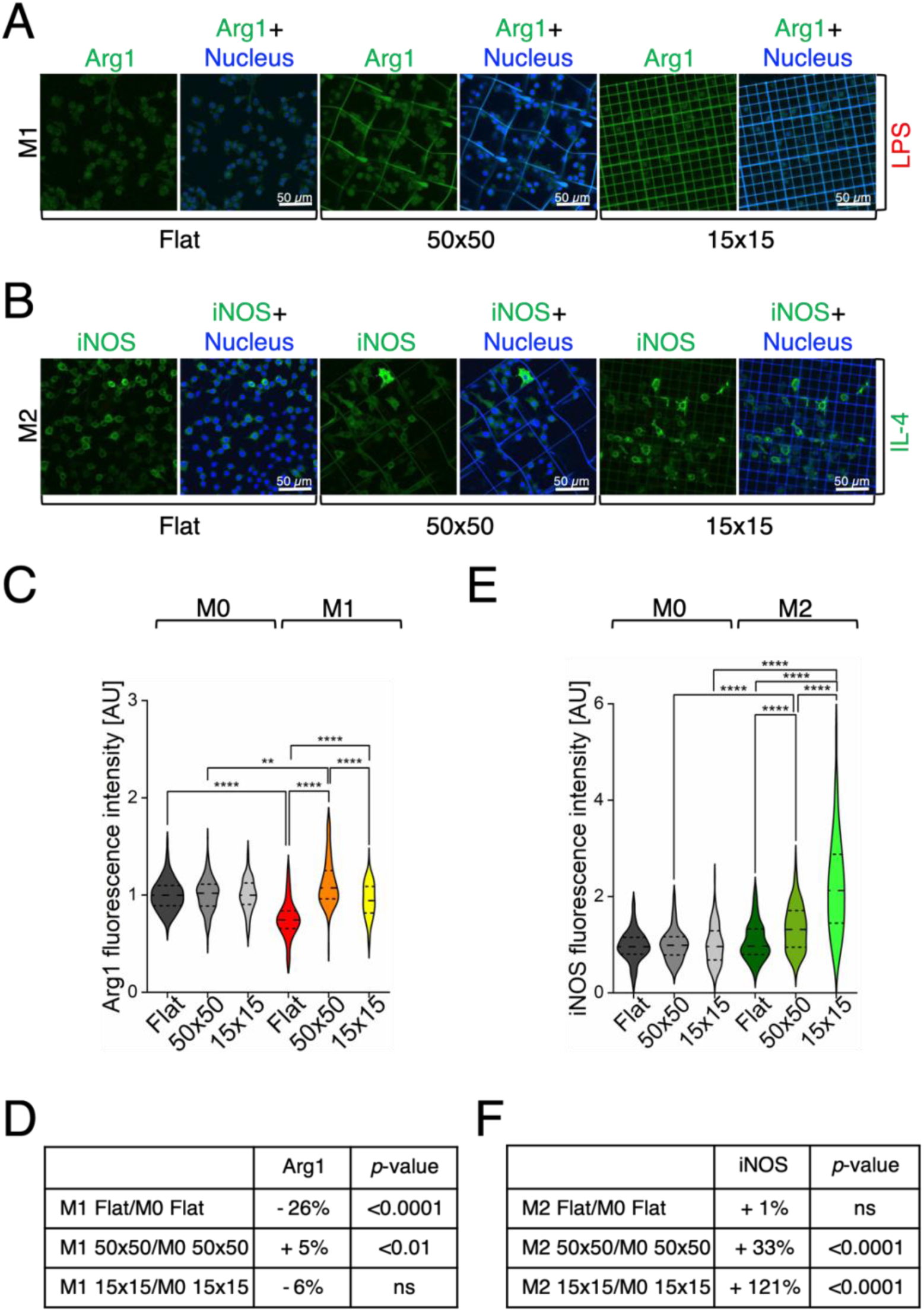
The architecture of two-photon polymerized microstructures can be harnessed to significantly modulate macrophage phenotype in an *in vivo*-like context. A-B) Representative confocal laser scanning microscopy images of LPS induced (M1) (**A**) and Il-4 induced (M2) (B) macrophages grown on Flat, 50x50, and 15x15 substrates, stained to visualize Arg1 (green, **A**), iNOS (green, **B**), and nuclei (blue). **C-D**) Quantification of Arg1 fluorescence intensity in M1 macrophages cultured on flat glass, 50x50, and 15x15 microstructures, normalized to the corresponding M0 controls (**C**). The variations (%) of iNOS expression are summarized in table (**D**). **E-F**) Quantification of iNOS fluorescence intensity in M2 macrophages cultured on flat glass, 50x50, and 15x15 microstructures, normalized to the corresponding M0 controls (**E**). The variations (%) of Arg1 expression are summarized in table (**F**).

## 4. Conclusion

Even though many studies have reported on the application of microstructured biomaterials to influence macrophage behaviour and to direct them towards pro-regenerative responses, a clear consensus on the ideal 3D geometry and pore size has not yet been reached. This research work presents an innovative strategy for modulating the activity and polarization phenotype of macrophages, thanks to the combination of both physical and chemical stimulations. The design and fabrication of our 3D fine microstructures by the two-photon polymerization technique allows to study the interaction with macrophages at the few micrometers scale, the closest to the cell’s size, and to investigate the synergistic effect of biochemical and physical cues, characteristic of the tissue microenvironment. Indeed, both the chemical signals and the 3D tissue architecture contribution to the behaviour and phenotype of macrophages were analysed. 3D microstructures were fabricated with two different porosities, 50x50x20 μm^3^ and 15x15x15 μm^3^ and, notably, both the morphology of the cells in terms of cytoskeletal organization, and the metabolic activity resulted to be significantly altered, suggesting a modulation in the mechanotransduction pathways. Interestingly, the 3D microstructures alone did not trigger macrophage polarization, underlining their suitability for *in vivo* use. Even more strikingly, scaffolds featuring 50x50x20 μm^3^ pores markedly increased Arg1 expression, indicating a shift toward an anti-inflammatory M2 phenotype, despite the presence of pro-inflammatory stimuli, thereby preserving the M1/M2 balance, essential for effective tissue regeneration. Conversely, microstructures with smaller pores (15x15x15 μm^3^) drove macrophages toward a pro-inflammatory phenotype, even in the presence of biochemical cues intended to promote tissue healing. Subsequently, the increased glycolytic activity observed in these macrophages suggests a metabolic environment less supportive of M2 polarization, which typically favours oxidative phosphorylation, highlighting the utility of these scaffolds as *in vitro* platforms for screening anti-inflammatory therapeutics and optimizing drug dosage regimens. This work is the first to our knowledge that completely avoids considering the contribution of the flat surfaces surrounding the fabricated 3D scaffolds, by analysing only macrophages effectively located inside the microstructures by means of imaging techniques. This represents a substantial advancement in the application of 3D architectures able to model the physiological environment upon implantation of a biomaterial and to modulate macrophage behaviour towards the desired phenotype. Indeed, the results obtained are crucial both for modelling pro-inflammatory conditions, and for guiding macrophage polarization towards pro-regenerative phenotype. This has direct implications for promoting tissue regeneration in conditions characterized by impaired healing, such as chronic wounds and ulcers. Furthermore, it supports the development of biomaterials decorated with 3D microstructures capable of modulating the foreign body response, thereby enhancing the integration of grafts and implants within the host tissue. These findings are pivotal for the tissue engineering community: they not only enable accurate modelling of acute inflammatory states but also provide a strategy to steer macrophage behaviour toward pro-regenerative profiles, paving the way for implants and grafts incorporating these microarchitectures to enhance tissue integration. Therefore, our results have immediate relevance in preclinical and clinical contexts, either to minimize acute inflammation and infection risk following device implantation, to encourage healing in chronic wounds or to shift the phenotype of macrophages in cancer immune therapy. Moving forward, we will focus on developing medical-grade resins to facilitate the translation of these insights into practical therapeutic applications.

### CRediT authorship contribution statement

CM and SC performed experiments. CM, SC and EJ acquired and analysed data. CM, SC, and EJ wrote the paper. GB supported *in vitro* experiments execution and scaffold fabrication. MV and CC supervised the process of scaffold fabrication. CM and EJ designed experiments and EJ supervised the study. GCe and RO developed the two-photon polymerization process and provided the laboratories to perform scaffold fabrication. GCh supported FLIM analysis interpretation. GCh, EJ, and MTR revised the manuscript. All authors contributed to this work and approved the submitted version.

## Acknowledgements

The authors thank Francesco Bonetto, alumnus of Politecnico di Milano, for his support in data collection, and Guangchen Wang, Ph.D. student from Trinity College of Dublin for technical assistance with Phyton coding.

## Appendix A. Supplementary data

Supplementary data to this article can be found online at [DOI].

## Research data availability

Data supporting the findings of this study are openly available in [repository name] at [URL or DOI]; the implementation of the phasor are available at [GitHub/repository link], ensuring full transparency and reproducibility.

## Declaration of AI-assisted technologies in the proofreading process

During the preparation of this work the authors used the free version of ChatGPT (GPT-3.5, OpenAI) in order to assist with English language proofreading of the manuscript. After using this tool, the authors reviewed and edited the content as needed and take full responsibility for the content of the published article.

## Appendix A. Supplementary data

**Supplementary Figure 1.**
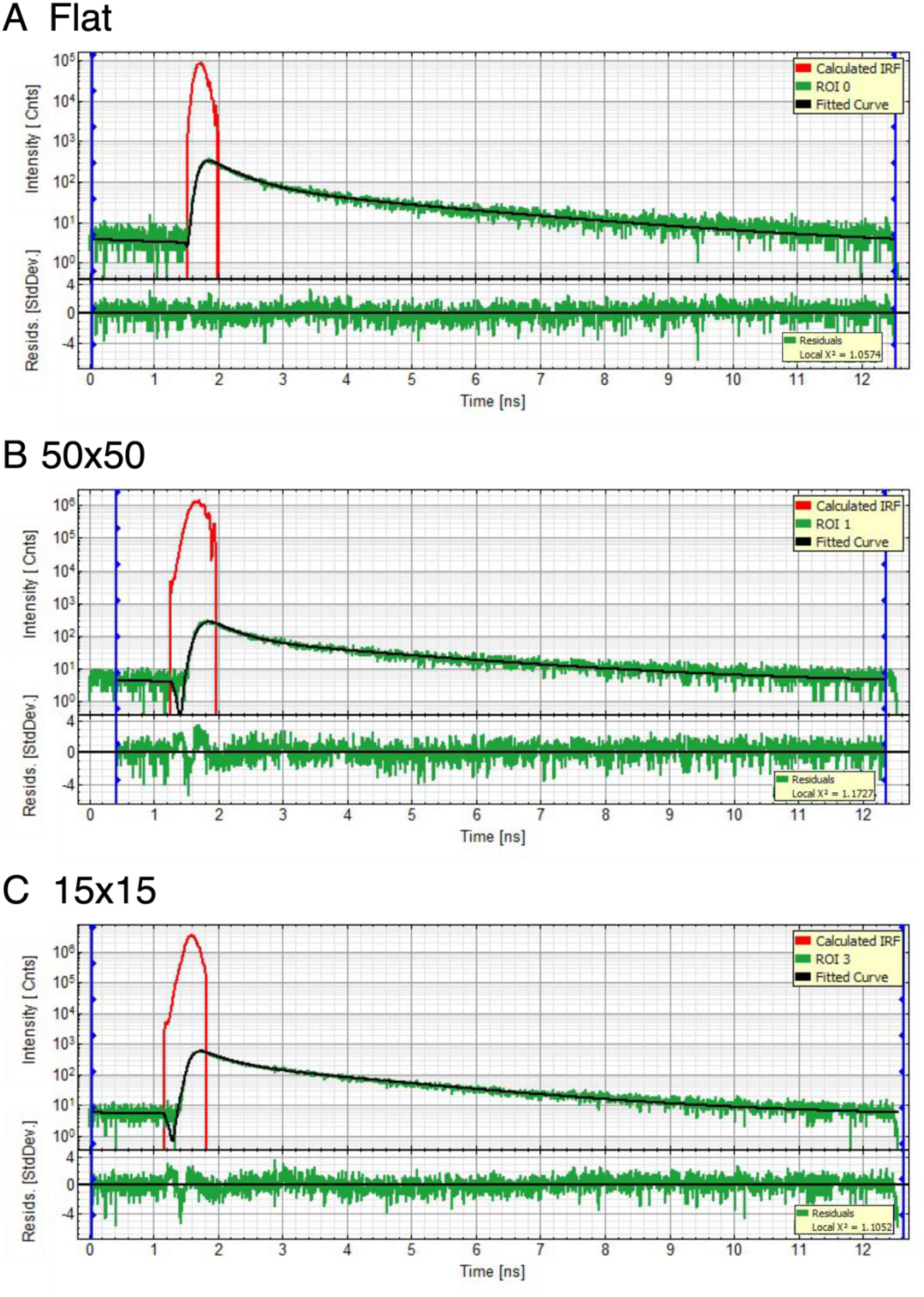
Representative fluorescence lifetime fitting curves. (**A-C**) Representative fluorescence decay curves from FLIM measurements acquired on Flat, 50×50, and 15×15 scaffolds, respectively. Graphs were exported directly from SymPhoTime 64, the software used for data analysis. In each plot, the green line represents the fluorescence intensity decay measured in a single manually drawn cellular ROI. The black line corresponds to the fitted decay model, while the red line shows the instrument response function (IRF). Below each decay curve, the residuals of the fit (green) are shown, along with the reduced chi-squared (χ²) values indicating the goodness of fit.

**Supplementary Figure 2.**
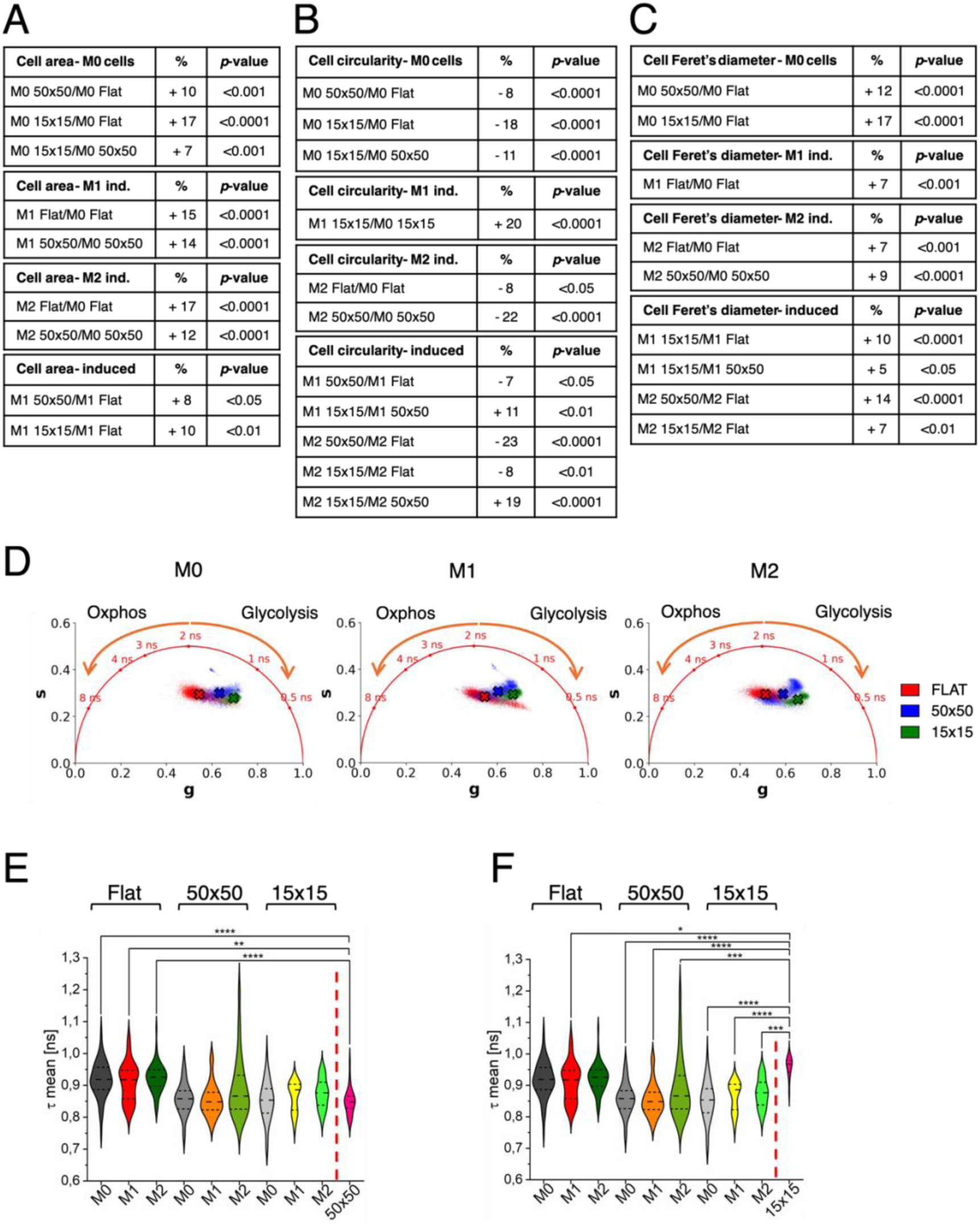
(**A-C**) Quantification of morphological parameters: cell area (**A**), circularity (**B**), and maximum Feret’s diameter (**C**) for M0, M1, and M2 macrophages cultured on Flat, 50x50, and 15x15 substrates in terms of variations (%). **D**) Comparison of metabolic activity across the same conditions, as assessed by phasor FLIM analysis. The crosses in the phasor plots represent the centroids of the data clouds, corresponding to the average fluorescence lifetimes; a shift toward the right (shorter lifetimes) indicates a more glycolytic metabolic state. **E-F**) Summary of metabolic variations in M0, M1, and M2 macrophages across the three substrates, alongside the mean fluorescence lifetime values for the two scaffolds: 50x50 (**E**) and 15x15 (**F**). Samples 50x50 and 15x15 on the *X* axis, separated by the dotted red line, indicate the 50x50 and the 15x15 empty scaffolds, respectively.

**Supplementary Video 1.**
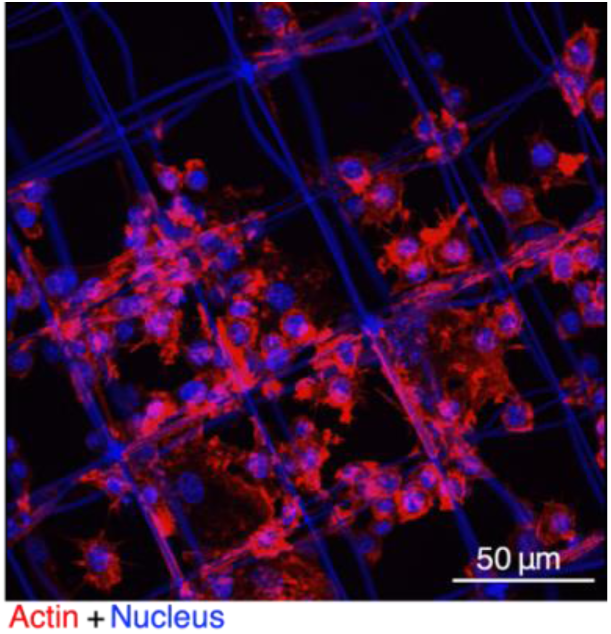
Representative 3D confocal laser scanning microscopy reconstructions of M0 macrophages cultured in the 50x50, stained for filamentous actin (red) and nuclei (blue). Due to the resin’s autofluorescence at the blue wavelengths, the scaffold is also visible in the blue channel alongside the cell nuclei.

**Supplementary Video 2.**
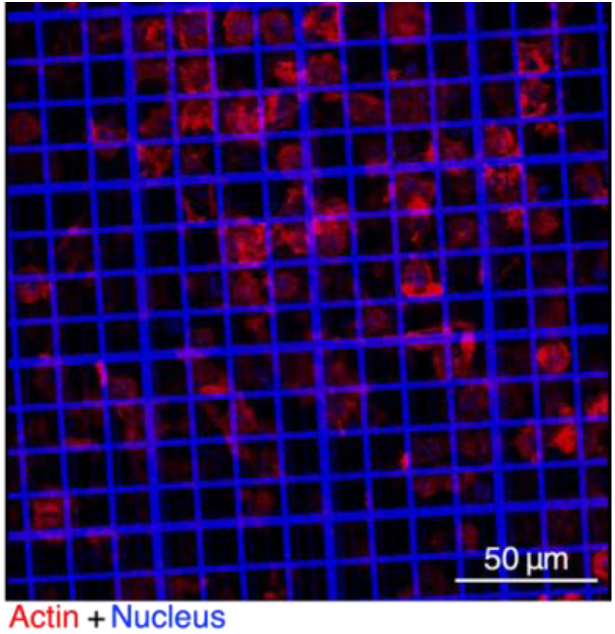
Representative 3D confocal laser scanning microscopy reconstructions of M0 macrophages cultured in the 15x15, stained for filamentous actin (red) and nuclei (blue). Due to the resin’s autofluorescence at the blue wavelengths, the scaffold is also visible in the blue channel alongside the cell nuclei.

**Supplementary Figure 3.**
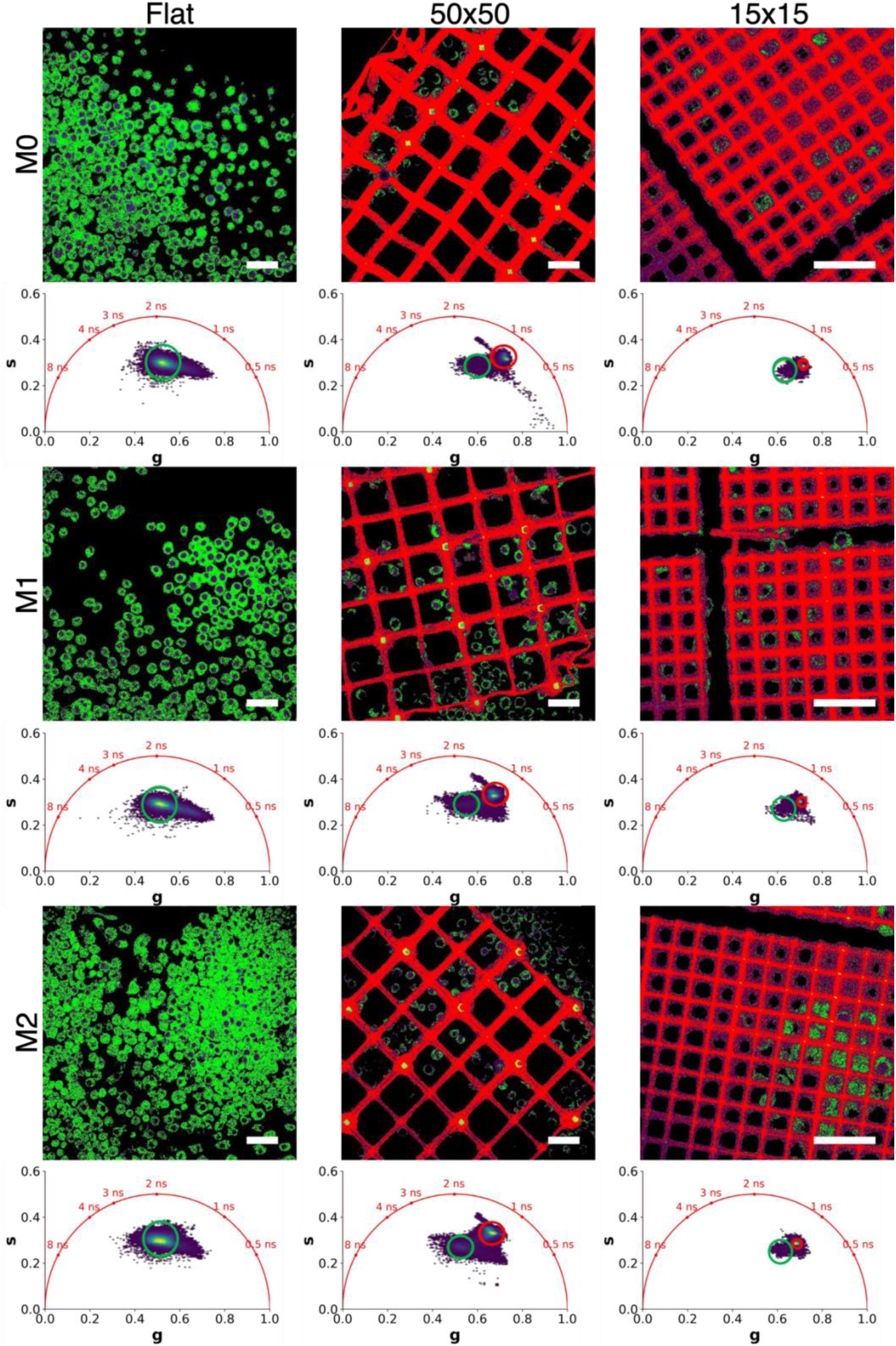
Representative fluorescence lifetime imaging microscopy (FLIM) images (top) and corresponding phasor plots (bottom) of M0, M1, M2 macrophages cultured on the flat glass substrate and in the 50x50 and 15x15 microstructures. Classification of cells (in green) and scaffolds (in red) performed using corresponding circles on the phasor plot. Scalebar 40 µm.

